# Bruch’s Membrane Contributes to the Structural Integrity of the Human Eye

**DOI:** 10.1101/2025.11.13.688375

**Authors:** Royston K.Y. Tan, Swati Sharma, Anita S.Y. Chan, Candice E.H. Ho, Fabian A. Braeu, Jost B. Jonas, Le Han, Xiaofei Wang, Yinling Zhu, Hwa Liang Leo, Martin L. Buist, Tin Aung, Shamira A. Perera, Michael J.A. Girard

**Affiliations:** Singapore Eye Research Institute, Singapore National Eye Centre, Singapore; Duke-NUS Medical School, Singapore; Singapore National Eye Centre, Singapore; Rothschild Foundation Hospital, Institut Français de Myopie, Paris, France; SERI-NTU Advanced Ocular Engineering (STANCE) Program, Singapore, Singapore; Key Laboratory for Biomechanics and Mechanobiology of Ministry of Education, Beijing Advanced Innovation Center for Biomedical Engineering, School of Biological Science and Medical Engineering, Beihang University, Beijing, China; Department of Biomedical Engineering, National University of Singapore, Singapore; Department of Ophthalmology, Yong Loo Lin School of Medicine, National University of Singapore and National University Health System, Singapore; Department of Biomedical Engineering, Georgia Institute of Technology and Emory University, Atlanta, United States; Department of Ophthalmology, Emory University School of Medicine, Atlanta, United States; Emory Empathetic AI for Health Institute, Emory University, Atlanta, GA, USA

**Keywords:** Bruch’s membrane, Sclera, Ocular biomechanics

## Abstract

**Purpose:** To investigate the contribution of the Bruch’s membrane and sclera tissues to the overall structural integrity of the ocular wall.

**Methods:** Twenty-three human globes were subjected to biomechanical testing. A piece of sclera measuring 5 x 5 mm was carefully removed at the nasal region, 2 mm away from the optic nerve head. The intraocular pressure was increased at approximately 1 mmHg/s until Bruch’s membrane-uvea-retina-tissue layer (BMUR) ruptured. Next, strips of sclera and Bruch’s membrane-choriocapillaris (BMC) complex were isolated from the superior fundus region. Uniaxial tension tests were performed at a strain rate of 0.01/s and sampling rate of 15 Hz. The tangent moduli of the BMC and sclera at 0.01, 0.02 and 0.03 strains were compared.

**Results:** The rupture pressure of the BMUR was 98.1 ± 21.4 mmHg. The tangent moduli of the BMC at 0.01, 0.02 and 0.03 strains were 2.96 ± 1.44 MPa, 7.68 ± 1.78 MPa and 9.43 ± 2.11 MPa, respectively, and the tangent moduli of the sclera at 0.01, 0.02 and 0.03 strains were 1.09 ± 0.80 MPa, 2.72 ± 1.67 MPa and 5.69 ± 3.27 MPa, respectively.

**Conclusion:** The BMUR was able to sustain relatively high IOP before rupturing. The uniaxial tensile tests showed that the BMC tangent moduli were about 3 times of those of the sclera at strains of 0.01 and 0.02. Although the sclera is approximately 47 times thicker, the BMC is still likely to make a significant contribution (3.51% to 7.42% at strain <0.03) to the overall structural strength of the ocular wall.

## Introduction

The human eye’s structural integrity is essential for optimal visual function, with Bruch’s membrane serving as a component of the ocular wall. Bruch’s membrane has a thickness of 2 – 4 µm and is located between the retinal pigment epithelium (RPE) and the choroid. According to generally accepted descriptions, it is comprised of five distinct layers: the RPE basal lamina, an inner collagenous layer, an elastic layer, an outer collagenous layer, and the basal lamina of the choriocapillaris.^1, 2^ Since the RPE with its basal membrane can detach from the inner collagenous layer (as is the case for retinal pigment epithelium detachments in eyes with age-related macular degeneration), the biomechanical intergrity of Bruch’s membrane may be reliant on just the inner collagenous layer, the elastic layer, and the outer collagenous layer. Bruch’s membrane facilitates nutrient and waste exchange between the RPE and choroid, provides structural support, and serves as an attachment site for the RPE.^3, 4^ During fetal development, Bruch’s membrane begins to form in the 7th week of gestation, prior to the development of the sclera, highlighting its foundational role in ocular development.^2, 5^ Its biomechanical properties were previously explored by Wang et al., and the study showed that the porcine Bruch’s membrane-choroid complex could sustain intraocular pressures of up to 82 mmHg before rupturing, indicating substantial structural resilience.^6^

Often viewed as the primary pressure-bearing structure of the eye, the sclera is undoubtedly one of the strongest ocular tissues. However, the eye may be anatomically differentiated into an inner globe, consisting of the uvea and all tissue central to the uvea (i.e., Bruch’s membrane, RPE, retina, vitreous body, and lens), and the sclera, covering the inner globe along with the cornea. The sclera is connected to the inner globe only at the scleral spur anteriorly and the peripapillary choroidal border tissue posteriorly. Consequently, the sclera can be peeled off the choroid outside of its adhesions at the scleral spur and the peripapillary choroidal border tissue. This leaves Bruch’s membrane as the strongest part of the inner eye globe. Bruch’s membrane also serves as a physical barrier between the retina and choroid,^3, 7^ and is capable of withstanding intraocular pressure (IOP). The sclera is only permeable to small solutes,^8^ and choroidal pressure remains relatively stable^9^ due to complex autoregulation between the intravitreal compartment (i.e., IOP) and choroidal blood flow.^10^ The biomechanics of a trans-Bruch’s membrane pressure transfer and pressure equilibrium in an elevated IOP state has remained an under-explored topic, made complex by the dynamic changes in choroidal volume.^11-13^ Thus, understanding the biomechanics of Bruch’s membrane and how IOP stresses the individual structural tissues could offer explanations for the development of various ocular conditions. In glaucoma, elevated IOP stresses ocular structures, including Bruch’s membrane, potentially altering its mechanical properties and influencing disease progression, in particular since the peripapillary end of Bruch’s membrane is connected through the peripapillary choroidal border tissue to the merging region of the lamina cribrosa with the peripapillary scleral flange. In our previous study, we conducted simulations demonstrating that increasing the stiffness of Bruch’s membrane by a factor of 13.5 resulting in an increase in prelamina strains. Although this effect was relatively small, it was more pronounced under elevated IOP.^6^ These findings suggest that Bruch’s membrane stiffness can influence the overall biomechanical response of the optic nerve to changes in IOP. Similarly, in pathological myopia, excessive axial elongation may impose abnormal strain on the ocular wall, affecting Bruch’s membrane’s integrity and leading to retinal complications. Staphyloma formation, characterized by scleral thinning and protrusion, may also be influenced by changes in Bruch’s membrane’s biomechanical properties, since it helps to maintain the structural stability of the sclera by anchoring the retina to the outer scleral layers and facilitating force transmission.^14^

Age-related changes in Bruch’s membrane, such as increased collagen cross-linking and lipid accumulation, might reduce its permeability, leading to waste product accumulation and complement system activation.^1, 7^ These alterations may disrupt the structural and functional roles of Bruch’s membrane, potentially contributing to conditions like age-related macular degeneration. Additionally, mutations in extracellular matrix proteins, such as fibulin-5, located within Bruch’s membrane, have been associated with drusen formation and cutis laxa,^15, 16^ further emphasizing its importance in ocular health.

The purpose of this research is to evaluate and estimate the biomechanical strength of the Bruch’s membrane-choriocapillaris (BMC) complex to access its contribution to the overall structural integrity of the ocular wall. We explored Bruch’s membrane together with the choriocapillaris, since it is difficult to remove the choriocapillaris from Bruch’s membrane with damage to the latter. Specifically, we aimed to perform biomechanical analyses by (1) performing a rupture pressure test on globes with a portion of the sclera removed, and (2) uniaxial tensile tests of the sclera and BMC to access the moduli of the tissues *ex vivo*.

## Methods

De-identified healthy human globes (*n* = 23) were purchased from Saving Sights (Kansas City, MO, USA), with a random globe extracted from separate individuals (11 male, 12 female, 10 left, 13 right, aged = 64.1 ± 12.8 years). The eyes were enucleated and placed in a moist chamber at approximately 0°C to 4°C within 12 hours post-mortem. The shipment was kept cold in ice and delivered to the Singapore Eye Research Institute (Singapore National Eye Centre, Singapore) within 72 hours post-enucleation.

### BMC pressure rupture experiment

A piece of sclera measuring 5 x 5 mm was carefully removed at the nasal region to avoid the macula, and 2 mm away from the optic nerve head (**Figure 1A**). Two 27G needles were inserted into the anterior chamber; one needle was connected to a pressure sensor (XP2i Digital Pressure Gauge, AMETEK Inc., Pennsylvania, USA) and the other needle was connected to a mobile water column to increase pressure. The IOP was increased at approximately 1 mmHg/s until the BMC ruptured (characterized by the peak pressure value prior to a sudden drop in reading), and the reading from the pressure gauge at the point of rupture was recorded.

**Figure 1.**
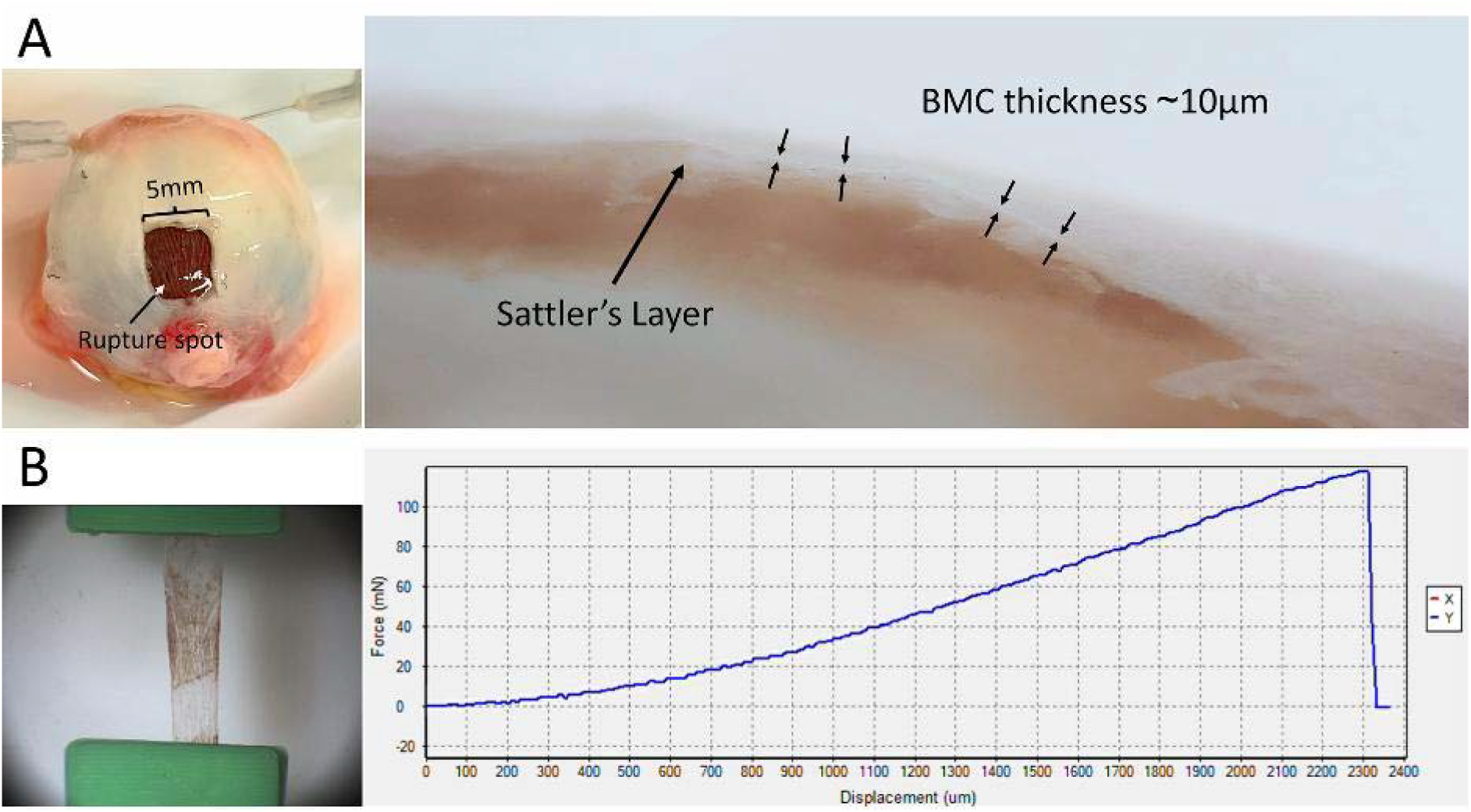
**(A)** The sclera was removed at the posterior nasal region for the rupture test (left), and subsequently the sclera and Bruch’s membrane-choriocapillaris (BMC) tissues (right) were isolated from the superior globe. **(B)** Example of the uniaxial tensile experiment performed on the BMC (24.0ºC, submerged in PBS, with preconditioning, sampled at 15 Hz at a strain rate of 0.01/s) to evaluate the tangent moduli at 0.01, 0.02 and 0.03 strains.

### Uniaxial tension of the Sclera and BMC

After the pressure test was completed, the globes were dissected. Strips of the sclera and BMC, running from the globe’s equator to the posterior pole, were isolated from the superior region. Bruch’s membrane strongly adhered to the choriocapillaris; hence the BMC was tested as a single unit. Using a combination of tying forceps (2-500-2E, Duckworth & Kent, Hertfordshire, UK), jewellers forceps (2-900E, Duckworth & Kent, Hertfordshire, UK) and Chihara conjunctival forceps (2-500-4E, Duckworth & Kent, Hertfordshire, UK), each BMC sample was prepared under a microscope to ensure proper removal of the retinal layers and the blood vessels from the Haller’s and Sattler’s layers. The resulting tissue was a thin transparent elastic layer (**Figure 1B and Figure 2**).

**Figure 2.**
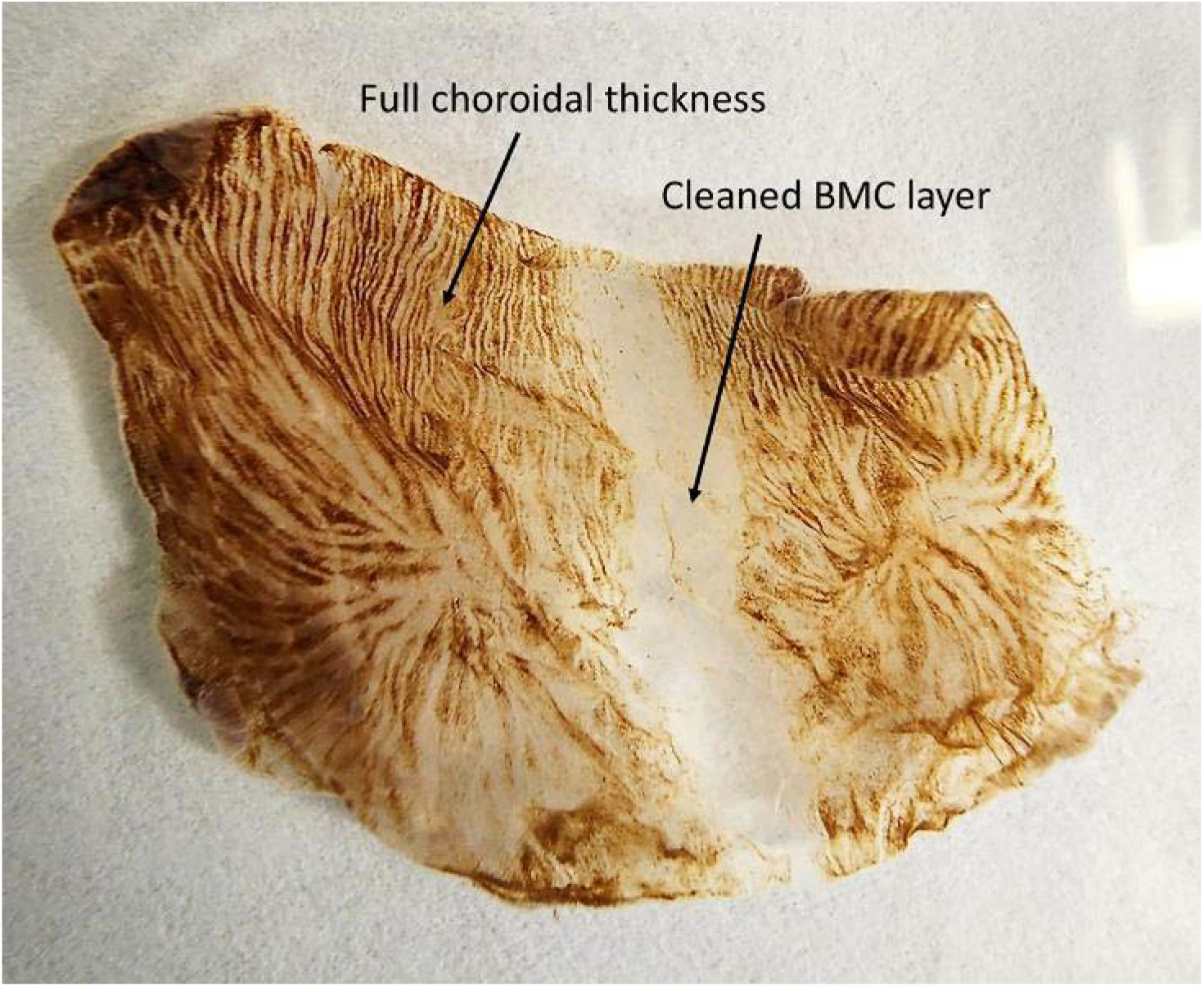
Sample preparation for the BMC involves removing the suprachoroid lamina, Haller’s layer and Sattler’s layer, leaving the choriocapillaris, which is strongly adhered to the Bruch’s membrane.

Optical coherence tomography (OCT) was used to determine the cross-sectional area of the sclera, and we measured the thickness of the BMC by microscopy (**Figure 3**). The technique to measure thickness depended on focusing the tissue sample at a specific plane. By folding the BMC sample into half, a portion of the transparent tissue thickness is oriented vertically at the folded edge (**Figure 3A and 3B**) for measurements. Since the tissue was very thin, OCT imaging was unable to accurately capture the thickness accurately. However, using a custom-designed OCT with a resolution of 1.55 µm/pixel, we were able to corroborate our microscopy results, which measured the BMC at approximately 7 – 13 µm and full choroid thickness at approximately 200 – 300 µm. Note that due to dehydration of the tissue under the microscope, we could not determine the thickness of each tested BMC sample. Hence, a separate set of BMC tissue samples were used for thickness determination and calculations, and the uniaxial tension data was processed with an approximated thickness of 10.26 µm for all samples.

**Figure 3.**
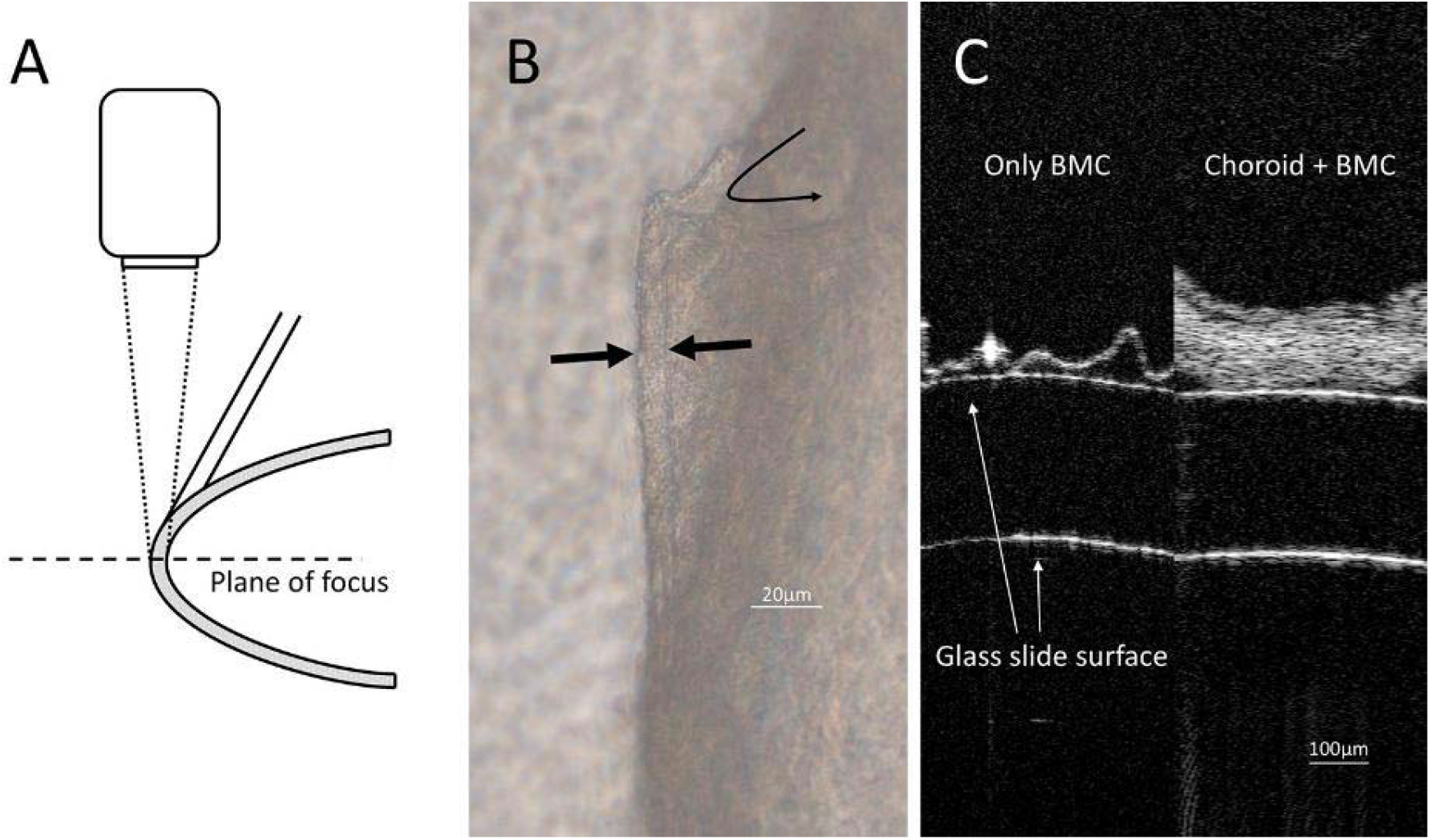
**(A)** Measuring the BMC thickness using microscopy by folding the tissue and adjusting the plane of focus. **(B)** By folding the tissue (curved arrow), we were able to obtain the focal point to measure the cross-section, which is the thickness of the tissue sample (straight arrows). **(C)** Using a custom high-resolution OCT, we were able to validate the thickness of the tissue to compare the BMC and full choroid thickness.

Preconditioning was performed on all samples at 0.05 strain for 10 cycles to achieve equilibrium, at laboratory temperature of 24.0°C with preloads of 10 mN and 1 mN for the sclera and BMC respectively. Uniaxial tension tests were performed on BMC and sclera samples at a strain rate of 0.01/s for 30 s and sampling rate of 15 Hz using the 0.5 N load cell from the BioTester 5000 (CellScale Biomaterials Testing, Waterloo, ON, Canada) (**Figure 1B**). The machine was calibrated as instructed in the user manual prior to every experimental session. A custom 3D-printed structure was designed to clamp the delicate tissue using sandpaper to prevent the tissue from tearing. A constitutive model for the entire data could have resulted in inaccurate curve fitting, thus only the local tangent moduli at low strains were determined. To calculate the tangent modulus of each sample, we used a third order polynomial fit (R^2^ > 0.99) to determine the local strain (ε) at ε < 0.05 and compared the BMC and sclera at 0.01, 0.02 and 0.03 strains (**Figure 4**), which are indicative of physiological strain levels. The results were reported as the mean ± standard deviation of all 23 samples.

**Figure 4.**
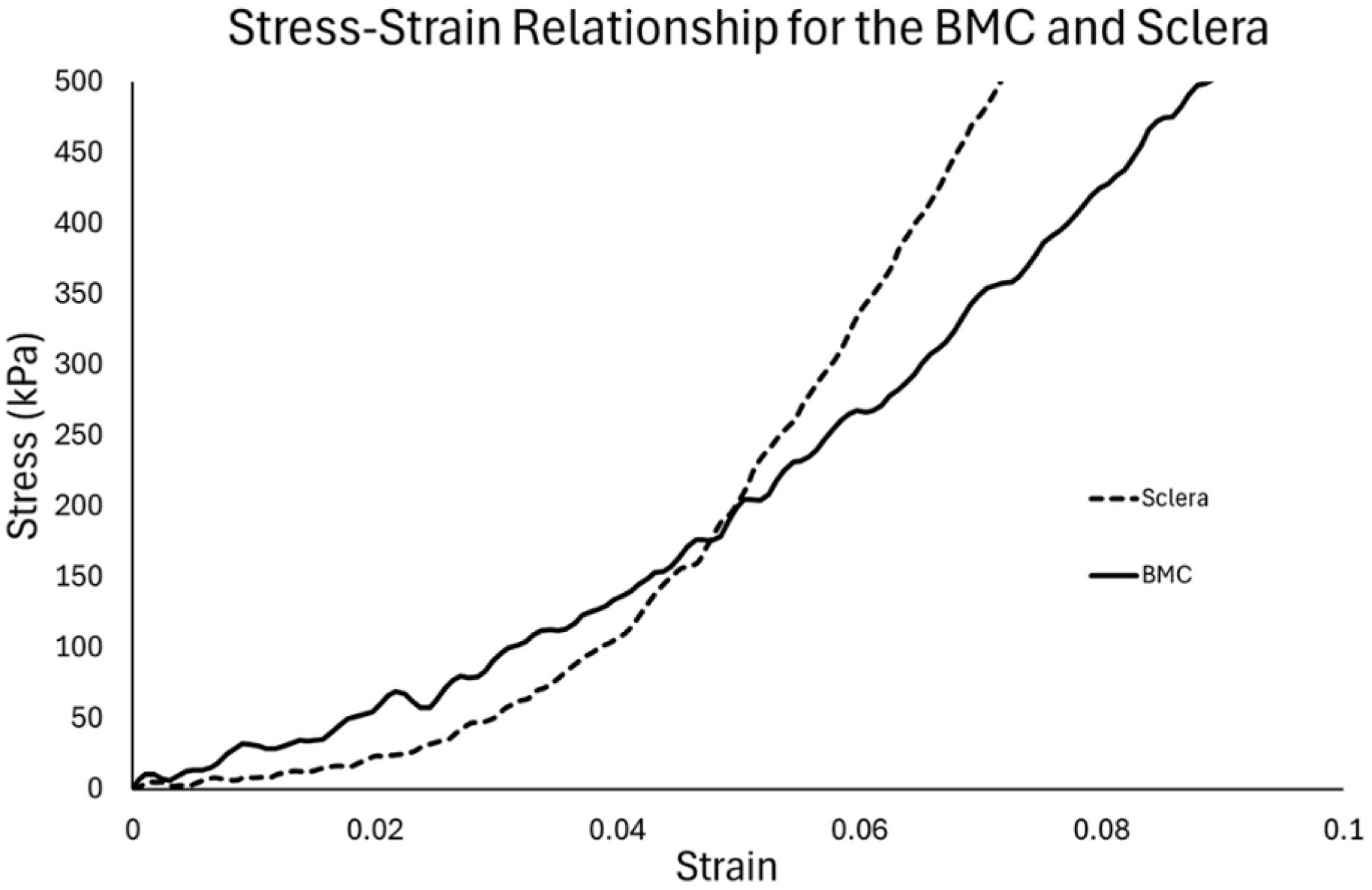
Examples of the stress-strain curves in this study. The data at low strains were reflective of in vivo conditions, and our results showed the BMC having a higher tangent modulus than the sclera below 0.03 strain.

### Histology

The BMC was isolated, with the RPE and choroidal blood vessels carefully removed manually, and subsequently fixed in 10% neutral buffered formalin solution (Leica Surgi path, Leica Biosystems Richmond, Inc.) for 24 hours. It was then dehydrated in increasing concentration of ethanol, cleared in xylene, and embedded in paraffin (Leica-Surgipath, Leica Biosystems Richmond, Inc.). Four-micron sections were cut with a rotary microtome (RM2255, Leica Biosystems Nussloch GmbH, Germany) and collected on POLYSINETM microscope glass slides (Gerhard Menzel, Thermo Fisher Scientific, Newington, CT). The sections were dried in an oven of 37°C for at least 24 hours. To prepare the sections for histopathological examination, the sections were heated on a 60°C heat plate, deparaffinized in xylene and rehydrated in decreasing concentration of ethanol. A standard staining procedure with hematoxylin and eosin (H&E) was performed.^17^ Staining using the Periodic Acid-Schiff (PAS) method (Leica Biosystems Richmond, Inc.) was also performed. A light microscope (Olympus BX42, Olympus Corporation, Tokyo, Japan) was used to examine the slides and images were captured using a digital pathology scanner (MoticEasyScan Pro 6, Motic, Xiamen, China).

## Results

### Pressure rupture and uniaxial tension experiments

The rupture pressure of the BMC was 98.05 ± 21.44 mmHg. After isolating the BMC, the remaining tissue thickness measured 10.26 ± 2.10 µm, whereas the sclera tissue thickness measured 486 ± 80 µm. The tangent moduli of the BMC at 0.01, 0.02 and 0.03 strains were 2.96 ± 1.44 MPa, 7.68 ± 1.78 MPa and 9.43 ± 2.11 MPa, respectively, and the tangent moduli of the sclera at 0.01, 0.02 and 0.03 strains were 1.09 ± 0.80 MPa, 2.72 ± 1.67 MPa and 5.69 ± 3.27 MPa, respectively.

### Histology

The microscope images affirmed the thickness measurements made using the folding technique on the fresh tissues (**Figure 5**). The H&E and PAS stains showed that the vessels from the Haller’s layer and Sattler’s layer were absent tissue, and our tissue isolation consisted primarily of the Bruch’s membrane, measuring approximately 2 – 3 µm, with the choriocapillaris beneath.

**Figure 5.**
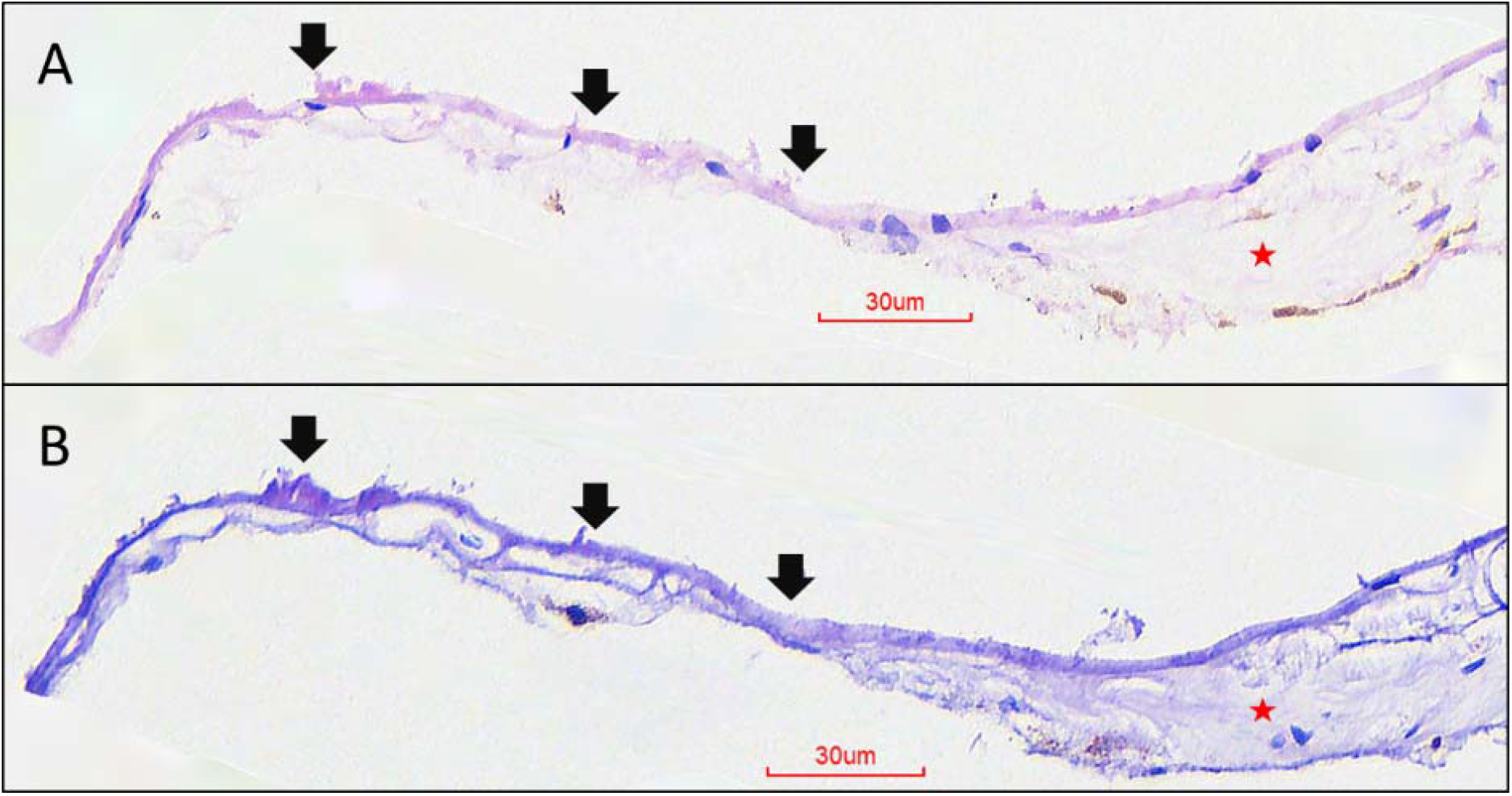
Histology results of the sample shown in Figure 2. **(A)** High power Hematoxylin and Eosin (H&E) stain showing the area of Bruch’s membrane without choroid (black arrows) compared to the adjacent pigmented areas with the choroid (starred). **(B)** High power Periodic Acid-Schiff (PAS) stain showing the area of Bruch’s membrane without choroid (black arrows) compared to the adjacent pigmented areas with the choroid (starred). The choriocapillaris remain firmly attached to the Bruch’s membrane.

## Discussion

We successfully isolated the BMC, by removing the adherent retina and the choroid except for the choriocapillaris. The thickness of our BMC measured using imaging was 9 – 11 µm, which was consistent with histological measurements^2^. This transparent layer was subjected to uniaxial tensile stretch tests to determine its biomechanical behaviour in comparison to the sclera. The globe pressure rupture test also provided corroboration of the BMC strength in containing high levels of IOP.

### The BMC was stiffer than the sclera and exhibited a more elastic behavior

The stress-strain behavior of pure collagen fibrils usually exhibits a linear response, with brittle failure (akin to a rubber band snap).^18^ It is the uncrimping of the uneven lengths and orientations of collagen fibers that results in its non-linear behavior. We observed the BMC to have a low non-linear behavior at the toe-region (at <0.05 strain), presumably due to the high composition of elastin offering a more linear response. Thus at these low strain levels, which represent physiological conditions, the BMC is stiffer than the sclera and able to resist more tissue stress. However, further stresses were resisted by the sclera when the level of deformation increased. It is important to note that the BMC may be stiffer, but the sclera still bears majority of the eye loads due to its greater thickness.

Differences in tissue composition between the sclera and BMC may explain the differences in biomechanical behavior. Bruch’s membrane consists primarily of collagen I, III and V, with a layer of elastin fibers, collagen VI and fibronectins within the elastic layer.^2^ The collagen IV at the two basal laminas may be of lower importance, since, as noted above, the RPE basal membrane can detach from Bruch’s membrane in various macula diseases. The choriocapillaris, which makes up between 83% and 58% of the BMC thickness and decreases with age,^19^ contributes substantially to the structure of the BMC. It is comprised primarily of collagen IV fibers forming a dense meshwork pattern.^20^ However, due to the spongy consistency of the choriocapillaris, the contribution of the choriocapillaris to the overall biomechanical strength of the BMC may be relatively low. Conversely, the sclera stroma mainly consists of bundles of collagen fibrils, forming thick lamellae up to 6 µm thick and aligned tangential to the scleral surface.^21^ Interfibrillar proteoglycans and glycoproteins such as decorin and biglycan surround a diffuse population of scleral cells,^22^ potentially facilitating mechano-transduction by regulating collagen fibril formation and organization.^23^ On the scleral outer surface, the episcleral tissue and Tenon’s capsule connect to the ocular muscles and function as supporting structures to transmit forces for eye movements. In short, the BMC is composed primarily of collagen fibers, with type IV being dominant,^24^ whereas the sclera contains bundles of collagen and elastin fibers with a rich extracellular matrix of fibroblasts and proteoglycans, and the differences in tissue composition may be one of the reasons for the differences in their respective stress-strain behaviors.

### The BMC is an important load bearing structure of the eye

It is possible to estimate the contribution of the BMC to the strength of the ocular wall (consisting of BMC and sclera combined) at specific strains. Under *in vivo* conditions, ocular tissues in human eyes experience strain levels between 0.01 to 0.09,^25, 26^ but are likely to be at the lower end of this range under physiological conditions. Once accounting for the thickness of the BMC (10.26 µm) and sclera (486 µm), it is possible to calculate, using the local tangent moduli, that the BMC bears 7.4%, 5.4%, 5.9% and 3.5% of the globe’s structural load at strains of 0, 0.1, 0.2 and 0.3, respectively. However, we believe that the autoregulation of choroidal pressure could create disproportionate load distributions. Our results show that the BMC can support up to 98 mmHg of pressure, whereas the sclera is able to sustain pressures up to 7275 mmHg.^27^ In a scenario where choroidal pressure is lower than intravitreal pressure, more of the IOP would be withstood by the BMC rather than the sclera (akin to an inflated water balloon inside a beach ball). In the event of a sharp impact, most of the force would be resisted by the sclera such that the BMC does not rupture.

### Myopia and staphylomas could be influenced by the BMC

Ocular deformations that lead to conditions such as myopia and staphylomas involve growth of the ocular wall, which includes both the BMC and sclera. Near distance lens accommodation involves contraction of the ciliary muscles pulling it anteriorly.^28^ Since the longitudinal ciliary muscle is connected to the basal membrane of the pigment epithelium in the pars plana region,^29^ and since that basal membrane continues into Bruch’s membrane, any accommodation-related forward traction of the ciliary muscle may lead to a strain on Bruch’s membrane, which is posteriorly fixed to the optic nerve head through peripapillary choroidal border tissue. In view of the relatively high biomechanical strength of Bruch’s membrane, as shown in the present investigation, one may discuss a biomechanical unit including Bruch’s membrane. This biomechanical unit starts at the corneal Descemet’s membrane, which is firmly through the transitional zone tissue and through the corneoscleral trabecular meshwork with the scleral spur^30^ which connects posteriorly with the longitudinal ciliary muscle. It is connected to the ciliary plana RPE basal membrane, which continues into Bruch’s membrane, and the posterior end of which connects through the peripapillary choroidal border tissue with the peripapillary border tissue of the peripapillary scleral flange. The latter border tissue crisscrosses with the collagen fibers from the peripapillary scleral flange which extend into the lamina cribrosa.

### Limitations

There were limitations in our experimental methods which may have affected the results. First, the average age of our samples was 64.1 years, which does not reflect the biomechanical behavior of a young adult’s tissues. The BMC is expected to be thicker and stiffer with increasing age,^7, 31, 32^ likewise the sclera would also have age-related changes.^33-35^ Second, the BMC thickness was microscopically determined using histologically processed tissue. Post-mortem tissue swelling and preparation-induced tissue shrinkage might have affected the tissue thickness. These factors however may affect vascularized tissue markedly more as compared to a structure consisting mostly of collagen and elastin like Bruch’s membrane. In addition, we used a separate set of samples to obtain an average thickness value and applied this value for our calculations. It meant that the BMC results were estimates unlike the sclera, which had the corresponding thickness measurements for each set of data. Finally, the pressure values for the globe burst experiment depended on the readings from the pressure sensor, which provided a 4 Hz output. Thus, our recorded data could have been lower than the actual pressure, since we needed the peak pressure prior to failure. It might have led to an underestimation of the burst pressure, and may only serve to corroborate the finding of a biomechanically relatively strong Bruch’s membrane.

## Conclusions

The BMC was able to sustain relatively high IOP before rupturing and the uniaxial tensile tests showed that the BMC tangent moduli were about 3 times of those of the sclera at strains of 0.01 and 0.02. It is possible that the mechanical forces within the eye are primarily buffered by the BMC at low strains and then transmitted to the sclera.

## Acknowledgements

The authors thank the donors of the National Glaucoma Research, a program of the BrightFocus Foundation, (G2021010S [MJAG]); the NMRC-LCG grant ‘TAckling & Reducing Glaucoma Blindness with Emerging Technologies (TARGET)’, award ID: MOH-OFLCG21jun-0003 [MJAG]; and the SERI-Lee Foundation grant (LF0622-03), for support of this research.

